# Reassessing Retinal Pigment Epithelial Ketogenesis: Enzymatic Assays for Ketone Body Levels Provide Inaccurate Results

**DOI:** 10.1101/2024.06.03.597221

**Authors:** Gillian A. Gulette, Daniel T. Hass, Kriti Pandey, Qitao Zhang, John Y.S. Han, Abbi Engel, Jennifer R. Chao, Nancy J. Philp, James B. Hurley, Jason M.L. Miller

## Abstract

The retinal pigment epithelium (RPE) is omnivorous and can utilize a wide range of substrates for oxidative phosphorylation. Certain tissues with high mitochondrial metabolic load are capable of ketogenesis, a biochemical pathway that consolidates acetyl-CoA into ketone bodies. Earlier work demonstrated that the RPE expresses the rate-limiting enzyme for ketogenesis, 3-hydroxy-3-methylglutaryl-CoA synthase 2 (HMGCS2), and that the RPE indeed produces ketone bodies, including beta-hydroxybutyrate (β-HB). Prior work, based on detecting β-HB via enzymatic assays, suggested that differentiated cultures of primary RPE preferentially export β-HB across the apical membrane. Here, we compare the accuracy of measuring β-HB by enzymatic assay kits to mass spectrometry analysis. We found that commercial kits lack the sensitivity to accurately measure the levels of β-HB in RPE cultures and are prone to artifact. Using mass spectrometry, we found that while RPE cultures secrete β-HB, they do so equally to both apical and basal sides. We also find RPE is capable of consuming β-HB as levels rise. Using isotopically labeled glucose, amino acid, and fatty acid tracers, we found that carbons from both fatty acids and ketogenic amino acids, but not from glucose, produce β-HB. Altogether, we substantiate β-HB secretion in RPE but find that the secretion is equal apically and basally, RPE β-HB can derive from ketogenic amino acids or fatty acids, and accurate β-HB assessment requires mass spectrometric analysis.

## Introduction

The retinal pigment epithelium and photoreceptors, collectively termed the “outer retina”, exist in a synergistic metabolic state (Adijanto et al., 2014; Bazan et al., 1994; Hurley, 2021; Kanow et al., 2017; Rodriguez de Turco et al., 1999; Sinha et al., 2020). Photoreceptor metabolism is characterized by the Warburg effect, in which aerobic glycolysis predominates despite ample mitochondrial mass. The oxygen and nutrients from the outer retina are derived from the choroidal blood supply and must transit the RPE. The accepted paradigm in the literature is that the RPE prefers fuels other than glucose, ‘sparing’ glucose for photoreceptors. In turn, the RPE relies on mitochondrial rather than glycolytic metabolism to support its anabolic and catabolic needs. It has been proposed that the RPE oxidizes fatty acids from the ingestion of lipid-rich photoreceptor outer segment (OS) tips and lipids from circulating lipoproteins. This lipid load can be metabolized via mitochondrial β-oxidation. The terminal step in β-oxidation produces acetyl-CoA, a substrate for the Krebs cycle and for ketogenesis. Ketone bodies are condensation products of acetyl-CoA and can be theoretically formed from any metabolite that produces acetyl-CoA, including fatty acids, certain amino acids, and glucose (Laffel, 1999). Historically, ketogenesis has been measured via β-HB levels, as β-HB tends to be produced in significantly higher abundance compared to other ketone bodies (Newman & Verdin, 2014). The RPE produces ketone bodies, specifically β-hydroxybutyrate (β-HB), and β-oxidation is thought to be a major contributor to RPE β-HB (Adijanto et al., 2014; Reyes-Reveles et al., 2017). RPE-derived β**-**HB has been reported to be preferentially released apically through monocarboxylate transporter isoform 1 (MCT1) (Adijanto et al., 2014). It has been postulated that photoreceptors have the capacity to uptake RPE-derived β-HB through the relevant MCT1/MCT7 transporters and utilize the ketone bodies for metabolic fuel (Adijanto et al., 2014).

Given the RPE’s capacity for ketogenesis and the potential utilization of ketone bodies by photoreceptors as a metabolic fuel, there has been interest in measuring β-HB as a proxy for RPE metabolic health and fatty acid metabolism (Adijanto & Philp, 2014). However, in this study, we found that enzymatic and mass spectrometric methods for detecting β-HB did not match. Indeed, an artifact in enzymatic assays arise when β-HB concentration is at the limits of detection. Using more sensitive mass spectrometry and isotopic tracing methods, we find that the RPE produces lower levels of β-HB than previously reported and transports the ketone bodies equally to the apical and basal sides. As β-HB levels rise, the RPE also begins consuming the metabolite. Finally, we find that RPE β-HB can be derived from both fatty acids and ketogenic amino acids, preventing the ketone body from being an exclusive read-out of β-oxidation. We suggest future RPE β-HB analysis be performed by mass spectrometry rather than enzymatic kits.

## Methods

Human pre-natal primary RPE cultures (hfRPE) were established according to previously published methods (Maminishkis et al., 2006; Zhang, Presswalla, Feathers, et al., 2019). Briefly, hfRPE were plated on microporous supports (Transwells; Corning 7200154) at Passage 1 and monitored for RPE-like features, such as cobblestone morphology, high pigmentation, and TEER of at least 500 Ω*cm2, before experimentation. Cultures were utilized approximately 2-4 months after plating on Transwells. HfRPE are regularly cultured in a complete α-MEM media (Maminishkis et al., 2006), with the addition of N1 Medium Supplement (Sigma Aldrich N6530), 1X GlutaMAX (Life Technologies Gibco 35050-061), Penicillin-Streptomycin (Life Technologies Gibco 15140-122), Taurine (Sigma T8691), Hydrocortisone-Cyclodextrin (Sigma H0396), 3,3’,5-Triiodo-L-Thyronine Sodium Salt (Sigma T5516), Non-Essential Amino Acid Solution (NEAA) (Life Technologies Gibco 11140-050), and 5% fetal bovine serum (FBS) (Atlanta Biologicals S11550H). This media will be referred to as RPE media hereafter.

Cells were incubated in RPE media (with or without FBS) or Earle’s Balanced Salt Solution (EBSS) with 200 μM palmitate and 1 mM carnitine in the apical and basal chamber of the Transwell, for anywhere from 2 to 48 hours. Palmitate utilized in this study was conjugated to fatty-acid free bovine serum albumin (BSA) at a ratio of 6:1, as previously published (Hass et al., 2023). For molecular weight size exclusion, conditioned media was placed in a 10 kDa molecular weight cut-off (MWCO) low-binding column (Microcon® MRCPRT010) and centrifuged at 14,000 x g for a total of 20-25 minutes, depending on loaded volume. The size-excluded flow-through was collected and run alongside the non-size-excluded samples using the AAT Bioquest Fluorometric Ketone Body kit (13831) or StanBio Colorimetric Ketone Body kit (SBHR100). The AAT Bioquest kit was run according to manufacturer instructions and fluorescence was measured at excitation= 540 nm, emission= 590 nm. The colorimetric StanBio kit was run exactly as previously described (Adijanto et al., 2014), recapitulating apical/basal media volume ratios (scaled to cell culture surface area), media composition, RPE culture type, and absorbance wavelength (492 nm). For all enzymatic assay kits, molecular weight size-excluded and non-size-excluded β-HB standard curves were created, ensuring that the MWCO columns were not sequestering β-HB. For each type of media incubated with hfRPE, the corresponding standard curve was composed of the particular media used for the culture. Importantly, FBS contains small amounts of β-HB, but this “background” β-HB was accounted for by employing the same percent FBS in the standard curves as percent FBS used for a given experiment. β-HB concentration in samples was determined using the linear regression equation of the respective standard curve.

For primary culture carbon tracing experiments, hfRPE was incubated in a mix of 5.5 mM glucose and 300 μM palmitate conjugated to BSA (Sigma A8806). In these experiments, either the glucose or palmitate was ^13^C-labeled, such that we incubated cells in 5.5 mM ^13^C_6_-glucose (Cambridge Isotope Laboratories Inc. CLM-1396-0) and 300 μM BSA-conjugated ^12^C-sodium palmitate (Sigma P9767), <u>or</u> ^12^C-glucose and 300 μM ^13^C_16_-sodium palmitate (Cambridge Isotope Laboratories Inc. CLM-6059-0). Medium from each treatment group was sampled at 0, 2, 4, 7, 10, and 13 hours after medium change. For mouse eyecup carbon tracing experiments, we cervically dislocated 3–5-month-old C57BL/6J mice, removed the eyes, and dissected RPE-choroid-sclera from each eye. Each tissue unit was incubated in 500 μL Krebs-ringer-bicarbonate (KRB) buffer/well of a 24-well plate. KRB was supplemented with 5 mM glucose, 100 μM palmitate, and 1 mM leucine. For each experimental group, the C^13^ label was either on glucose (^13^C_6_) or leucine (^13^C_5_). An hour after the beginning of the incubation, the tissue was flash frozen in liquid N_2_. Tissue was homogenized and metabolites were extracted in 80% methanol, supplemented with a 10 μM methyl succinate internal standard.

Liquid chromatography-mass spectrometry (LC-MS) was performed by the Michigan Regional Comprehensive Metabolomics Resource Core (MRCMRC) (Ann Arbor, MI, USA.). Metabolites were extracted from samples at 1:5 in extraction solvent (80:20 methanol/water) along with 1 μM 3-Hydroxybutyrate(D2) and 1 μM 2-Hydroxybutyrate(D3) internal standards. Samples were reconstituted in 100 μl methanol: water (8:2 by volume). Metabolites were analyzed on an Agilent system consisting of an Infinity Lab II UPLC coupled with a 6545 QTof mass spectrometer (Agilent Technologies, Santa Clara, CA) using a JetStream ESI source in negative mode. Data was processed using MassHunter v B.07.00 (Agilent Technologies, Santa Clara, CA). Absolute quantification, when displayed, is based off a multi-point standard calibration.

Gas chromatography-mass spectrometry (GC-MS) was performed at the University of Washington. Metabolites were extracted from cell culture supernatant in an extraction buffer consisting of 80% MeOH, 20% H2O, and 10 μM methyl succinate (Sigma, M81209). The extraction buffer was equilibrated on dry ice, then 150 μl was added to each sample. The supernatant was then dried under vacuum and stored at -80°C until derivatization. Dried samples were derivatized by adding 10 μl of 20 mg/mL methoxyamine HCl (Sigma, 226904) dissolved in pyridine (Sigma, 270970) incubating at 37°C for 90 minutes, adding 10 μl tert-butyldimethylsilyl-N-methyltrifluoroacetamide (Sigma, 394882) and incubating at 70°C for 60 minutes. 1 μl of derivatized samples were injected into an Agilent 7890/5975C GC-MS system. Selected-ion monitoring was used to determine metabolite abundance, as previously described (Du et al., 2015). Peaks were integrated in MSD ChemStation E.02.01.1177 (Agilent Technologies), and correction for natural isotope abundance was performed with IsoCor, v1.0 (32). Corrected metabolite signals were converted to molar amounts by comparing metabolite peak abundances in samples with those in a homemade mixture of metabolite standards. Known amounts of standards were extracted, derivatized, and run alongside samples in each experiment. The resulting data was used to generate a standard curve that allowed for metabolite quantification.

## Results

Previous reports have used enzymatic assays to quantify β-HB secretion for RPE. These assays rely on the conversion of β-HB to acetoacetate by recombinant 3-hydroxybutyrate dehydrogenase. Oxidation of β-HB is coupled to the reduction of nicotinamide adenine dinucleotide (NAD+), which forms nicotinamide adenine dinucleotide hydride (NADH). For the StanBio colorimetric assay, NADH then reacts with an added tetrazolium salt, INT, in the presence of diaphorase to produce color that is detected by absorbance (Adijanto et al., 2014; Reyes-Reveles et al., 2017). For the AAT Bioquest fluorometric assay, a proprietary fluorescent NADH sensor is implemented.

Initially, we were interested in measuring β-HB secretion as a proxy for RPE β-oxidation utilizing more physiologic/complete media than the EBSS media previously employed for RPE ketogenesis assays. When we incubated hfRPE in complete media and assayed for β-HB secretion with an enzymatic kit, we saw high tonic levels of RPE β-HB secretion. However, we were unable to see an increase in β-HB secretion when we added a small molecule that increases β-oxidation in hfRPE, the acetyl-CoA carboxylase inhibitor firsocostat (Figure 1A). In contrast, when assessed by mass spectrometry, firsocostat increased β-HB levels (Figure 1B). To reconcile this discrepancy, we characterized two enzymatic assay kits used in prior RPE research further: the AAT Bioquest Fluorometric Ketone Body kit (13831) (Zhang et al., 2021; Zhang, Presswalla, Calton, et al., 2019) and the StanBio Colorimetric Ketone Body kit (SBHR100) (Adijanto et al., 2014; Miyagishima et al., 2021; Reyes-Reveles et al., 2017).

**Figure 1:**
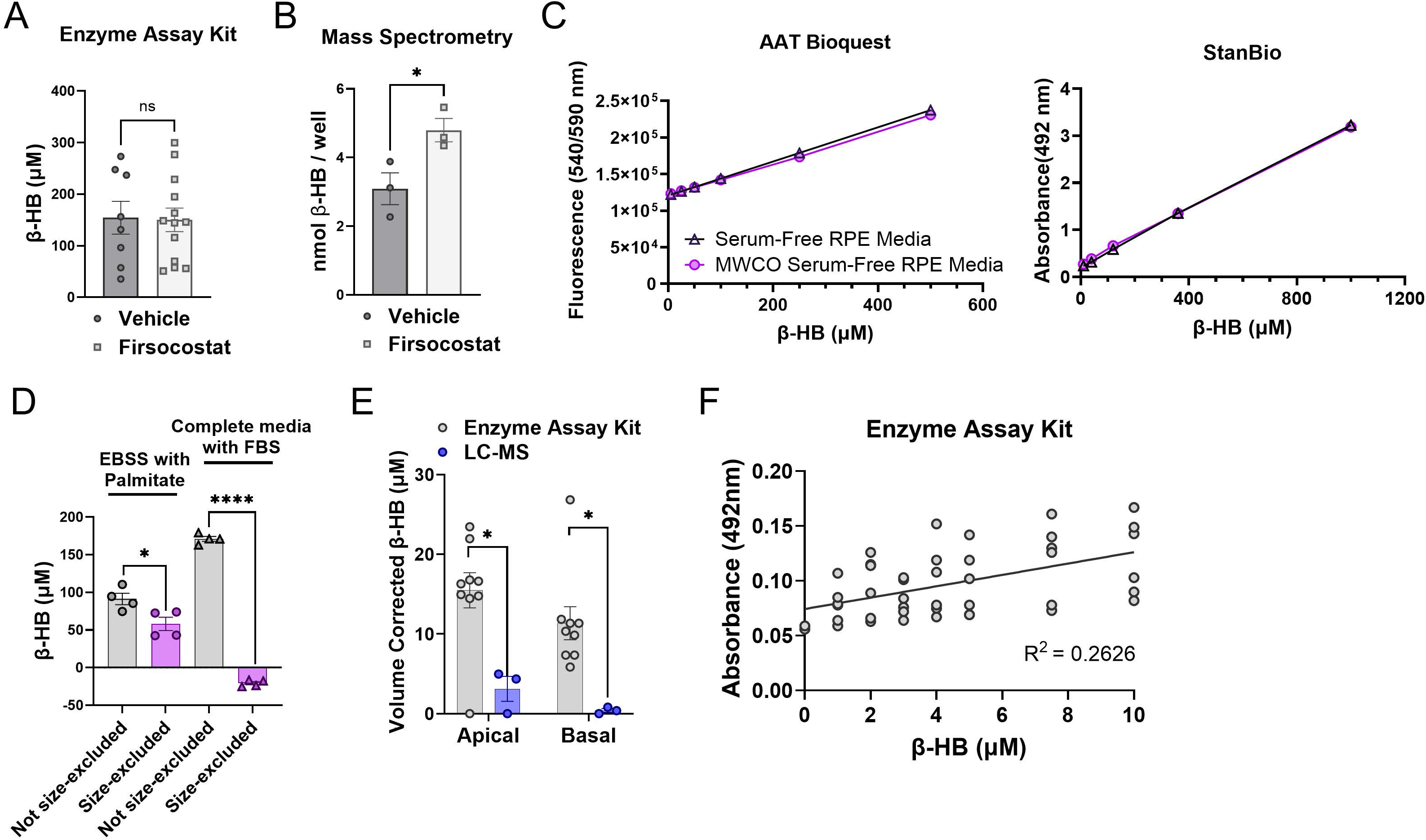
β-HB secreted by the RPE in culture cannot be accurately measured by enzymatic assay kits. (A) Firsocostat, a small molecule known to increase RPE β-oxidation, does not lead to increased β-HB production when measured via an enzymatic assay kit. Lipid-rich media was applied to hfRPE with 1 μM firsocostat or vehicle control. Apical conditioned media was collected, and β-HB was measured using the AAT Bioquest β-HB assay kit (t-test, ns, p=0.9153). (B) Firsocostat increases β-HB production when measured by mass spectrometry. Lipid-rich media was applied to hfRPE with 100 nM firsocostat or vehicle control. β-HB abundance in apical media was determined using GC-MS and quantity of β-HB was extrapolated from standards of known concentration (t-test, *, p<0.05). (C) Size-exclusion of media via MWCO columns does not remove β-HB. Utilizing the AAT BioQuest kit (left) or StanBio kit (right), along with serum-free standard RPE media, standard curves were assembled ranging from 0-1000 μM β-HB. (D) Size-exclusion of RPE supernatants via MWCO columns causes dramatic differences in measured β-HB concentration. Media was incubated over hfRPE for 3–24 hours and then the apical chamber supernatant was collected and measured by enzymatic assay kit. (t-test, *, p<0.05; ****, p<0.0001). (E) Comparison of β-HB concentrations from the same RPE supernatant samples utilizing enzymatic assay versus quantitative LC-MS demonstrates wide discrepancies. RPE supernatants incubated with EBSS + 200μM palmitate + 1 mM carnitine and collected after 3 hours incubation were measured by the StanBio enzymatic β-HB assay kit or via LC-MS employing deuterated internal standards as well as a multi-point standard curve. Experimental conditions were designed to exactly replicate RPE β-HB experiments reported previously. Apical and basal β-HB concentrations are mathematically corrected for the differences in media volume in the apical versus basal chambers (150 μL apically, 450 μL basally) (t-test, *, p<0.05). (F) The β-HB enzymatic assay kit used in key previous publications of RPE β-HB secretion is not sensitive enough to accurately measure β-HB at the concentrations secreted in RPE culture. LC-MS of RPE supernatants in (E) reveals β-HB concentrations below 10 μM. Technical replicates of β-HB concentration standards between 0-10 μM in EBSS supplemented with 1 mM carnitine and 300 μM palmitate was measured by the StanBio β-HB. When fit with a linear curve, R^2^ = 0.2626. All error bars for Figure 1 are S.E.M.

We first characterized the assays by determining whether a large molecular weight component of the RPE supernatant may interfere with accurate β-HB quantification. For example, the presence of NADH in secreted exosomes by the RPE could lead to a false positive signal (Arslan et al., 2013; Wang et al., 2020). We utilized enzymatic assays on RPE supernatants that had or had not been filtered through a 10 kDa MWCO column. To ensure the size-exclusion columns do not sequester β-HB, we created standard curves using serum-free RPE media with standards ranging from 0-1000 μM β-HB. The β-HB standards were measured by fluorometric or colorimetric enzymatic assay either as is, or after they were filtered through the MWCO column. When comparing the standard curves derived from media with and without size-exclusion, the MWCO column did not affect the apparent concentrations of β-HB standards (Figure 1C). We then used these assays to determine β-HB content in supernatant from cells cultured for 4-48 hours. MWCO filtered supernatant contained a lower β-HB concentration than samples not subjected to size exclusion (Figure 1D).

Given conflicting β-HB values depending on whether a size-exclusion column was utilized, we compared quantification of the same RPE supernatant by colorimetric assay to quantification by LC-MS/MS. Similar to the previously published method (Adijanto et al., 2014) for quantifying RPE β-HB, we incubated mature hfRPE on Transwells in media containing 5mM glucose in EBSS with 200 μM BSA-conjugated palmitate and 1 mM carnitine for 3 hours. The ratio of apical to basal media volumes and apical media volume to Transwell cell surface area were both identical to prior publications (Adijanto et al., 2014; Reyes-Reveles et al., 2017). The enzymatic assay, performed without size exclusion, measured an apparent apical β-HB concentration of 15.47 μM and an apparent basal β-HB concentration of 11.35 μM. In contrast, LC-MS assessment of the same sample resulted in an apical concentration of 3.11 μM and basal concentration of 0.82 μM (Figure 1E).

Given these low β-HB concentrations, we re-examined the enzymatic assay’s ability to reliably detect β-HB from 0-10 μM. Using technical replicates of the standards, the enzymatic assay at these lower concentrations results in a curve that yielded an R^2^ value of ∼0.26. This spread between technical replicates suggests that the enzymatic assay is not sensitive enough to distinguish differences in RPE β-HB secretion under the experimental conditions previously published (Figure 1F).

With an understanding that enzymatic assays will not accurately assess a low level of RPE β-HB, we returned to mass spectrometry to gain a fuller understanding of RPE ketogenesis. All LC-MS included deuterated internal standards and multi-point standard curves to enable full quantitative analysis. We first measured β-HB in the supernatants of RPE cultures exposed to RPE media with various concentrations of serum for 48 hours. Regardless of serum concentrations, we saw a similar amount of β-HB production (Figure 2A). Because serum itself contains β-HB, media *not* exposed to cells had increasing amounts of β-HB at higher serum concentrations (Figure 2A). The “cap” on β-HB concentrations despite increasing amounts of lipid and β-HB available in media with higher serum levels suggests that β-HB may also be consumed by RPE and that by 48 hours, a steady state between production and consumption is achieved. Indeed, when we trace β-HB concentrations in media over time, we see an initial peak and then decline, consistent with production then consumption (Figure 2B).

**Figure 2:**
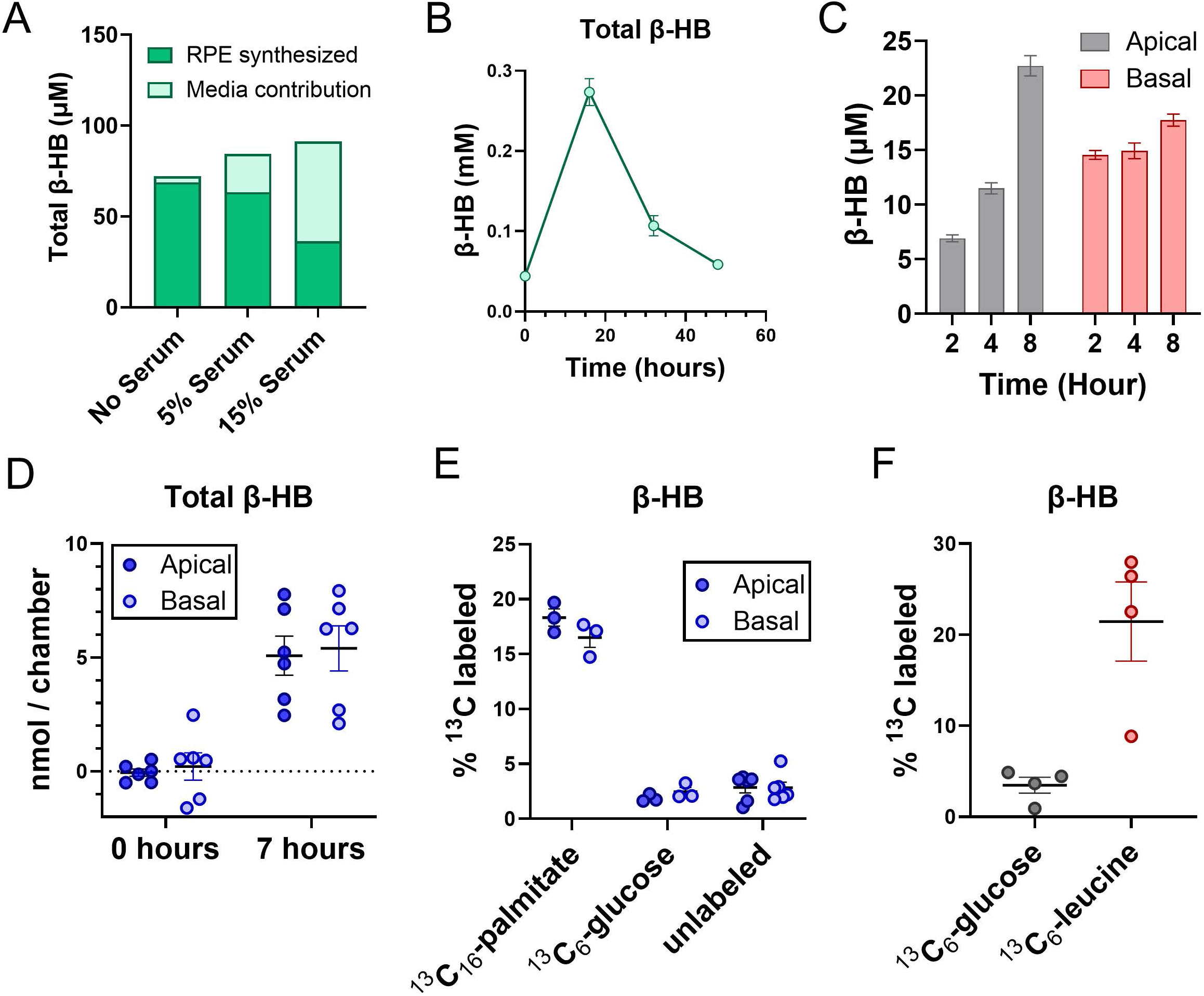
RPE β-HB is generated and secreted equally to the apical and basal chambers, consumed as RPE nutrient status declines, and made by both fatty acid and amino acid metabolism but not glycolysis. (A) β-HB secretion by hfRPE, as assessed by quantitative mass spectrometry, reaches a steady-state at 48 hours, regardless of how much starting β-HB (from FBS) is in the media. Medium that was collected after 48-hour cell exposure (RPE synthesized) and media with no cell exposure (media contribution) was quantitatively measured by LC-MS. (B) An equilibrium between β-HB production and consumption is reached by 48 hours. Conditioned RPE media supplemented with 300μM palmitate was collected over 48 hours and β-HB was quantitatively measured by GC-MS. Initially, a peak in β-HB concentration is seen at 16 hours, but the subsequent decrease in β-HB suggests the RPE begins consuming the β-HB. (C) The RPE has a steady increase in apical and basal β-HB secretion over shorter periods of time. hfRPE cultures in lipid-rich RPE media, with β-HB measured via quantitative LC-MS. (D) The RPE releases β-HB equally to the apical and basal chamber after 7 hours as measured by GC-MS. (E) RPE β-HB derives from fatty acid much more than glucose metabolism. Either palmitate or glucose was C^13^ labeled in otherwise identical standard serum-free RPE media. hfRPE was incubated with C^13^ labeled media for 7 hours, collected from apical and basal chambers, and C^13^-labeled β-HB was assessed by GC-MS. (F) A substantial portion of RPE β-HB also derives from ketogenic amino acids. Either leucine or glucose was C^13^ labeled in KRB buffer containing glucose, palmitate, and leucine. Mouse eyecups were incubated for 1 hour in such media and transfer of C^13^ label to β-HB was monitored after a 1-hour incubation. All error bars in Figure 2 are S.E.M

While evidence supports a peak in β-HB production at 16 hours, we sought to understand β-HB dynamics at earlier timepoints, similar to the timepoints employed for our colorimetric enzymatic assays. As seen in Figure 2C, RPE cultures incubated with RPE media containing lipids for 2, 4, and 8 hours demonstrate a steady increase in secreted β-HB both apically and basally over time.

To determine the degree of apical vs. basal RPE β-HB secretion and determine how strongly different metabolic substrates contribute to RPE β-HB production, we turned to ^13^C tracing experiments. We incubated RPE cultures with identical serum-free RPE media containing standard glucose concentrations (5.5 mM) and BSA-conjugated palmitate (300 μM) but varied whether the ^13^C label was on palmitate or glucose. In contrast to previous reports suggesting RPE β-HB is predominately secreted apically (towards photoreceptors), we found β-HB secreted equally to the apical and basal chambers (Figure 2D). We also found, in contrast to palmitate, that glucose has negligible contribution to RPE-secreted β-HB apically or basolaterally (Figure 2E). To determine if other metabolites can also contribute to β-HB production in the RPE, we incubated mouse eyecups in EBSS media with 5 mM glucose, 100 μM palmitate, and 1 mM leucine. In separate samples, the ^13^C label was on glucose or leucine, but media compositions were otherwise identical. While ^13^C-glucose again failed to label β-HB, labeling of the ketone body from ^13^C-leucine was robust (Figure 2F). This suggests that β-HB production derives from at least two sources in the RPE – amino acids and fatty acids.

## Discussion

The RPE faces a large daily lipid load, including lipid-rich OS phagocytosed from photoreceptors and lipoprotein uptake from the choroidal circulation. It has been widely accepted that a significant portion of this lipid load is metabolized via RPE β-oxidation, producing energy for the RPE and decreasing its reliance on glucose, which may then be spared for photoreceptor consumption. The end-product of β-oxidation is acetyl-CoA, which feedback inhibits further β-oxidation and glycolysis. To reduce acetyl-CoA levels and potentially provide photoreceptors with an additional energy source, the RPE was hypothesized to be capable of undergoing ketogenesis, with secretion of the ketone bodies preferentially to the apical side where they could be taken up by photoreceptors as fuel.

In our efforts to understand RPE β-oxidation, we utilized enzymatic detection of the dominant RPE ketone body, β-HB, as a proxy. Enzymatic assays have been the leading method by which RPE ketogenesis has previously been explored. Here, we report that the enzymatic kits are not sensitive enough to measure β-HB at the levels produced and transported out of RPE cultures. A contaminating factor is secreted by RPE and increases the apparent β-HB signal detected by the enzymatic kit. This contaminating factor has a sufficiently high molecular weight such that it is removed by a 10kDa size-exclusion column. Future studies assessing RPE ketogenesis should therefore utilize using mass spectrometry or similar size exclusion columns.

A reassessment of RPE ketogenesis utilizing mass spectrometry in this study reveals that the RPE transports β-HB to both the apical and basal sides, in contrast to previous reports (Adijanto et al., 2014) suggesting apically dominated secretion. The concentration of β-HB in the RPE extracellular space increases initially, but then peaks and declines, suggesting the RPE is capable of both producing β-HB in times of nutrient excess and consuming β-HB as nutrients are depleted. Therefore, for experiments strictly focused on ketogenesis and not β-HB utilization, RPE supernatant needs to be assayed early after media change (within ∼16 hours). Finally, our analysis demonstrates that RPE β-HB can derive from both fatty acid and ketogenic amino acid metabolism, but not glucose metabolism, making β-HB an inaccurate marker by itself for assessing β-oxidation.

## Acknowledgements

All experiments adhered to the approved IACUC protocol for the Hurley lab at the University of Washington and the ARVO statement for the use of animals in ophthalmology research. HFT research received review and approval or exemption from IRB at University of Washington and University of Michigan.

